# Evolutionary Isolation and Phylogenetic Diversity Loss Under Random Extinction Events

**DOI:** 10.1101/143594

**Authors:** Mike Steel, Vahab Pourfaraj, Abhishek Chaudhary, Arne Mooers

## Abstract

The extinction of species at the present leads to the loss of ‘phylogenetic diversity’ (PD) from the evolutionary tree in which these species lie. Prior to extinction, the total PD present can be divided up among the species in various ways using measures of evolutionary isolation (such as ‘fair proportion’ and ‘equal splits’). However, the loss of PD when certain combinations of species become extinct can be either larger or smaller than the cumulative loss of the isolation values associated with the extinct species. In this paper, we show that for trees generated under neutral evolutionary models, the loss of PD under a null model of random extinction at the present can be predicted from the loss of the cumulative isolation values, by applying a non-linear transformation that is independent of the tree. Moreover, the error in the prediction provably converges to zero as the size of the tree grows, with simulations showing good agreement even for moderate sized trees (*n* = 64).

## 1. Introduction

Biodiversity is generally defined as the ‘variety of life’ [7], and much effort is expended to minimise its loss in the face of anthropogenic activity [15, 26]. One conceptually straightforward measure of the biodiversity encompassed by a subset (e.g. by the bird species found in a wildlife preserve) is the evolutionary history that the subset embodies [27]. This is usually operationalized as Faith’s Phylogenetic Diversity (PD) [2] or the sum of the edge lengths of the minimum spanning tree that connects a subset of the leaves (here, the species) with each other and with the the root of some larger tree (e.g. the phylogeny of all bird species). This measure scales reasonably with the size of the subset, is set-monotonic and submodular [23], and attributes more biodiversity to subsets of distantly related leaves over subsets of more closely related leaves. The original article that introduced the measure [2] has been cited nearly 1300 times^1^, and academic applications of PD have been well-received (see, e.g. [4]). However, PD has never, to our knowledge, been explicitly implemented in a conservation intervention, such as allocating resources to sets of species that encompass more rather than less PD.

Because every leaf of a phylogenetic tree will contribute a measurable amount of PD to a defined subset, leaf-specific diversity measures are possible: the simplest is just the length of the pendant edge leading to that leaf, or the decrement in PD if that leaf is lost. So, for example, species on longer pendant edges are more isolated on the tree, contribute more PD to the tree, and therefore may warrant especial conservation attention [14]. This concept of the ‘evolutionary isolation’ value of species was extended to several *ad hoc* measures by Redding [18, 19], specifically his ‘fair proportion’ and ‘equal splits’ measures (defined in the next section).

The fair proportion measure has been the focus of several high profile papers advocating the use of evolutionary history for conservation (see, e.g. [5, 11, 25]), and, importantly, is the basis of the ‘Edge of Existence’ conservation programme (see [10] and edgeofexistence.org). Species-specific measures need to be calculated only once and, once generated, can be used in a variety of settings without specialist phylogenetic expertise. However, they have also been criticised by Faith [3], who showed with specific examples how preferentially conserving leaves that score high for these isolation measures may not minimize loss of evolutionary history. Given the continued use of evolutionary isolation, we return to this question here and show how, under simple models of diversification and extinction, the summed ‘fair proportion’ or ‘equal splits’ values of the species that are lost from a tree through extinction strongly predicts the concomitant loss of PD.

### 1.1 PD and isolation indices

Let *T* be a rooted binary phylogenetic tree with branches *e* of lengths λ_*e*_. Given a subset *Y* of the leaf set *X* of *T*, let *PD*(*T,Y*) be the phylogenetic diversity of *Y* on *T* (i.e. the sum of the lengths of the branches connecting the leaves in *Y* and the root of *T*), and let *PD*(*T*) be the total length of *T* (the sum of all branch lengths; thus *PD*(*T*) = *PD*(*T,X*)). Let *φ* = *φ_T_* be an isolation index (e.g. fair proportion or equal splits), which satisfies:

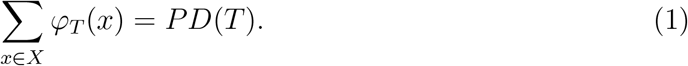

The *fair proportion* isolation index for leaf *x* is:

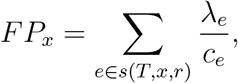

where *s*(*T, x, r*) is the set of edges in the tree *T* between leaf *x* and the root *r*, λ_*e*_ is the length of edge *e*, and *c_e_* is the size of the subclade each edge *e* in s subtends, such that every edge is divided uniquely among the leaves, satisfying Eqn. (1) above.

The *equal splits* measure also uniquely apportions each internal edge on the path from a leaf to the root to that leaf, but rather than splitting an edge ‘fairly’ among all the leaves in the clade it defines, it apportions each edge fairly to the two (or more) sister clades it defines. This means that a leaf gets an exponentially decreasing portion of the internal edge lengths as a function of the number of splits between it and the edge:

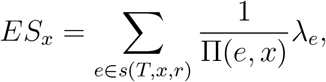

where Π(*e,x*) = 1 if *e* is a pendant edge incident with *x*; otherwise, if *e* = (*u,v*), then Π(*e*, *x*) is the product of the out-degrees of the interior vertices of *T* on the directed path from *v* to leaf *x*. For example, if *T* is binary and there are *k* interior edges separating *e* and *x*, then Π(*e,x*) = 2^*k*^. Essentially, the length of each edge is evenly distributed at each branching point (regardless of how many leaves are in each subtree).

The fair proportion index has been shown to be formally equivalent to a leaf’s contribution to the PD of a random sized random subset (the Shapley Index, [9], [23]). Though less studied, the equal splits index (on binary trees) can be conceptualized as the expected contribution of a leaf to the PD remaining if each of the subclades branching off the path from the leaf to the root were to independently become extinct with probability 0.5.

We will use *Y* to denote the leaves that survive some extinction event at the present and *E* = *X* – *Y* to denote the leaves that become extinct. For a subset *E* of *X*, let:

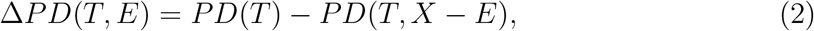

which is the loss in PD if the species in *E* become extinct. For a subset *Y* of *X*, and an isolation index *φ_T_*, let 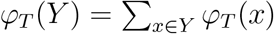 and let:

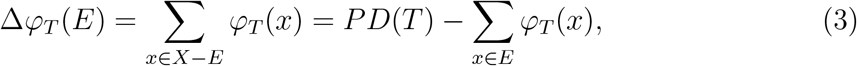

which is how much the sum of the *φ_T_* indices decreases if the species in *E* become extinct.

### 1.2. Convergence of diversity indices on birth—death trees under a ‘field of bullets’ model

Consider a phylogenetic tree generated by a constant rate birth–death process, with speciation and extinction rates *b* and *d* respectively. We will assume throughout that *b* > *d*. Let us grow this tree for time *t* and consider the resulting reconstructed tree *T_t_* (i.e. the tree based on the tips that survive to the present) with leaf set *X_t_*. Let us now select each leaf of *T_t_* independently with probability *p* (these survive), let *Y_t_* be the set of these surviving species and let *E_t_* = *X_t_* – *Y_t_* be the species that become extinct. This is the simple ‘field of bullets’ model of extinction at the present [17], operating on the reconstructed tree generated by a constant rate birth–death model. Note that there are two random processes at play here: firstly the process that generates the tree, and then the process that prunes leaves at the present.

### 1.3. Main result

Let *S_t_*(*p*) be the random variable corresponding to the PD of the randomly selected set of leaves (each chosen independently with probability *p*) on the random (reconstructed) tree *T_t_*.

Note that *S_t_*(1) = *PD*(*T_t_*) is the total length (the sum of the edge lengths) of *T_t_* and Δ*PD_T_t__* (*p*) := *S_t_*(1) – *S_t_*(*p*) is the loss of PD. If *E_t_* is the set of species that become extinct then, in terms of the earlier notation (Eqn. (2)), we can write Δ*PD_Tt_* (*p*) = Δ*PD*(*T_t_*, *E_t_*).

Let Δ*φ_T_t__*(*p*) = Δ*φ_T_t__*(*E_t_*) which (by Eqn. (3)) is the random variable that measures the loss in the sum of the isolation indices of the leaves of the random (reconstructed) tree *T_t_* when the species in the random set *E_t_* become extinct.

#### Theorem 1.

(i) *For a Yule pure-birth model,*

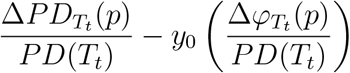 *converges in probability to 0 as t* → ∞, *where* 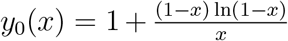. *Moreover, the slope of the curves y* = *y*_0_(*x*) *is* 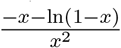 *for* 0 < *x* < 1, *which converges to* 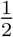 *as x approaches* 0 *from above*.
(ii) *For a birth-death model with b* > *d* > 0, *let θ* = *d*/*b*, *and let*

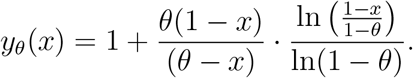 *Conditional on the reconstructed tree having at least one leaf*, 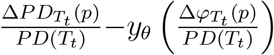 *converges in probability to 0 as t* → ∞. *Moreover, for d* > 0, *the slope of the curves y* = *y*_0_(*x*) *as x approaches* 0 *from above is given by* 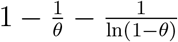.

**Remarks:** Stated less formally, Theorem 1 can be written as:

> ‘Proportion of PD lost = *y_θ_*(proportion of isolation index lost)’

Of course the terms on each side of this ‘equation’ are random variables, and Theorem 1 asserts that the informal equation becomes exact (i.e. the difference between the two expressions in the informal equation tends to zero) as *t* becomes large (or, more generally, if λ*t* becomes large). The curves described in Theorem 1 are illustrated in Fig. 1.

**Figure 1.**
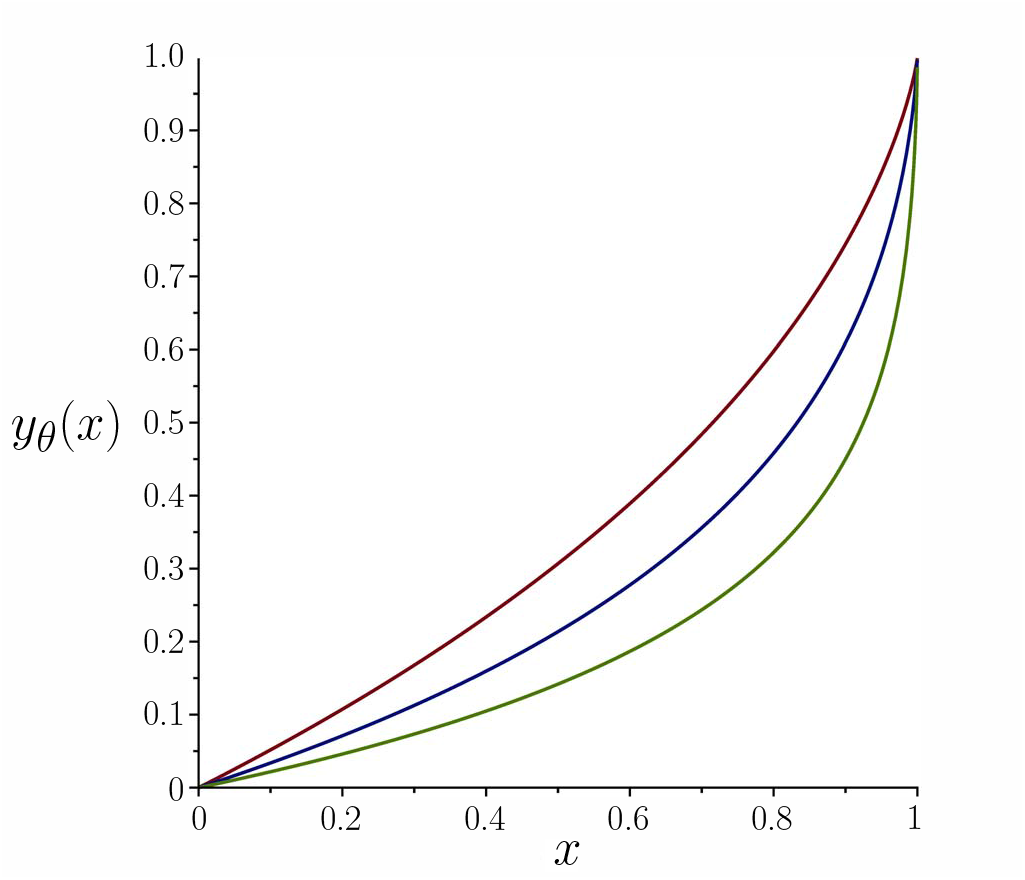
The curves of *y* = *y_θ_*(*x*) describing how the proportion of PD loss is related to the proportion of isolation score loss (in the limit of large *t*), for pure-birth where *d* = 0 (top curve), birth-death with *θ* = *d*/*b* = 0.9 (middle curve), and birth-death with *θ* = *d*/*b* = 0.99 (bottom curve).

Simulations, as presented in Fig. 2, show that Theorem 1 provides a reasonably unbiased prediction even for very moderate sized trees (*n* = 64).

Notice also that the function 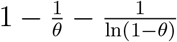 in Part (ii) of Theorem 1 is undefined at *θ* = 0 (i.e. for pure-birth trees); however, its limit as *θ* → 0+ exists and agrees with the value 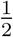 in Part (i). The curve for this function is illustrated in Fig. 3.

*Proof of Theorem 1*. The proof relies on two lemmas.

For the first of these, let

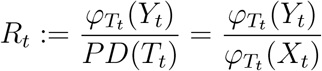

be the random variable which measures the proportion of the total isolation index that is spanned by the surviving species (under a field of bullets model with survival probability *p*) on *T_t_* (and for *d* > 0 conditional on there being at least one species in the reconstructed tree prior to the application of the field of bullets model).

#### Lemma 1.

*For both the fair proportion and the equal splits isolation indices, the random variable R_t_ converges in probability to p as t* → ∞.

*Proof*. By definition,

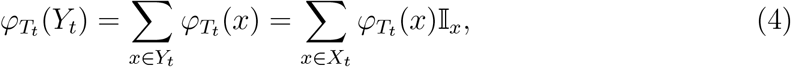

where 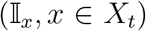 are independent and identical Bernoulli random variables that take the value 1 with probability *p* and zero otherwise. Thus we have the following equation for the conditional expectation of *R_t_* given *T_t_*:

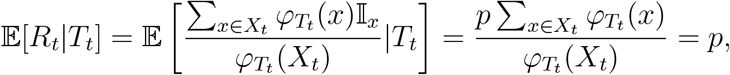

where the last equation follows by the linearity of (conditional) expectation, and Eqn. (1) (which holds for both fair proportion and equal splits) and Eqn. (4), noting that 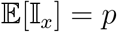 (note also that by conditioning on *T_t_* the term in the denominator of the fraction inside the expectation is treated as a constant). Thus, by the law of total expectation, we have 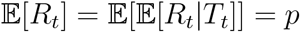.

**Figure 2.**
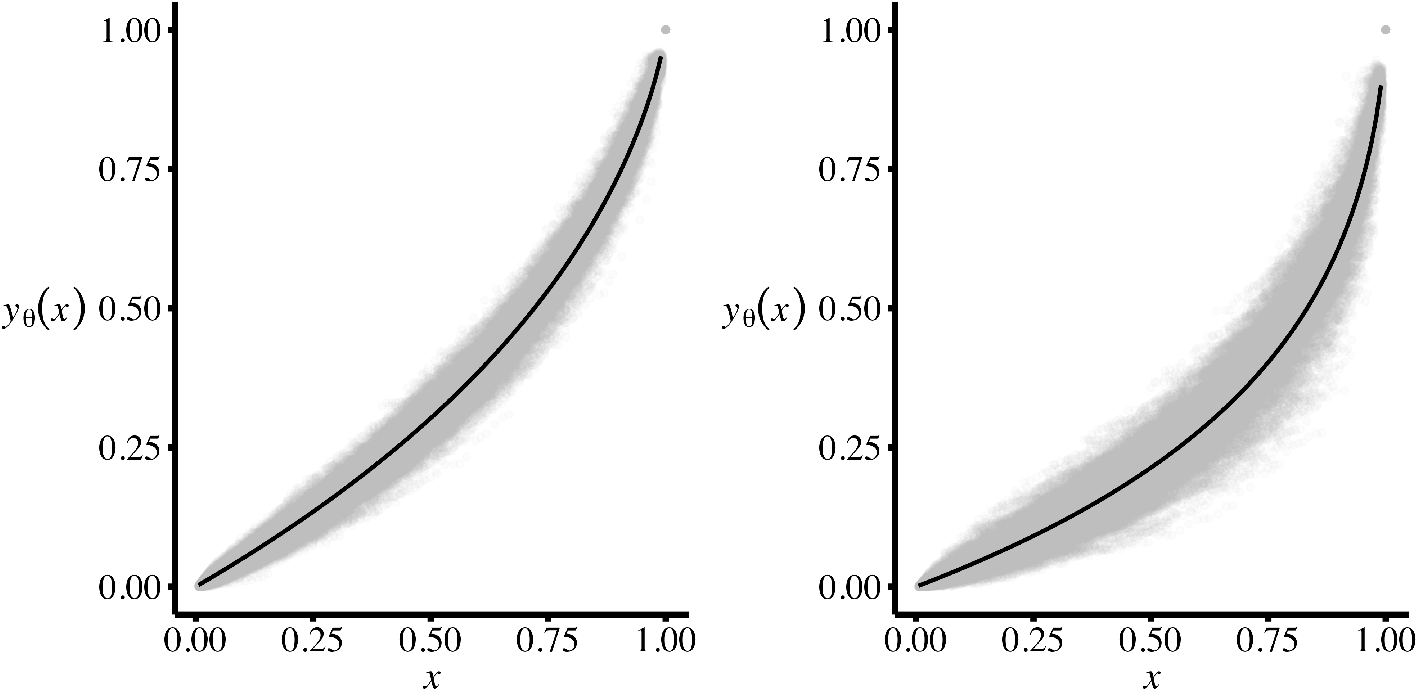
The loss of phylogenetic diversity versus the loss of cumulative fair proportion on simulated trees. The left panel shows the relationship across 1000 birth-death trees of size 64, with *b*=1 and *d*=0.1. The right panel shows the relationship across 1000 trees of size 64, with *b*=1 and *d* = 0.9. The expected relationship from Theorem 1 is depicted as the solid black line.

**Figure 3.**
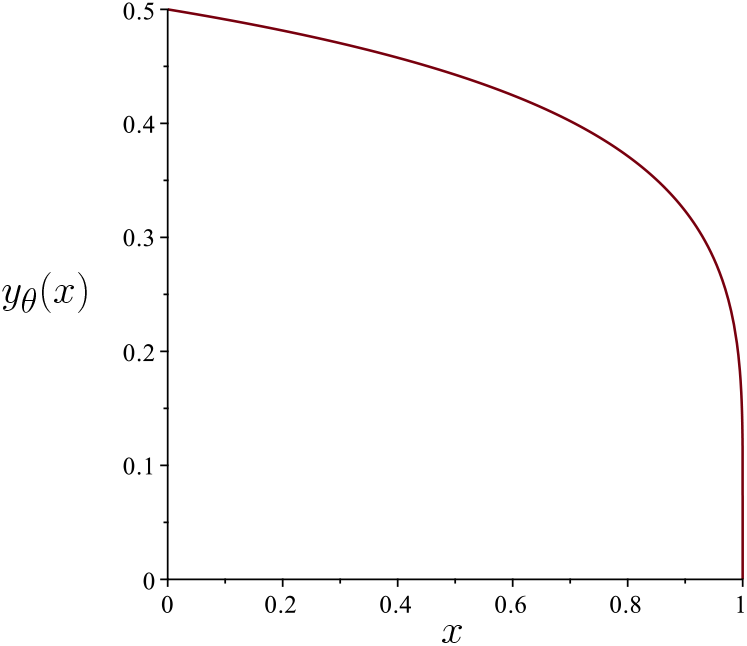
The ‘initial slope’ function 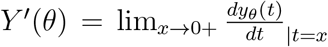 versus *θ* = *d*/*b* ∈ [0,1]. *Y*’(*θ*) is the slope of the function *y_θ_*(*x*) in the limit as *x* tends to zero from above, given in Theorem 1 (Part (i) for *θ* = 0 and Part (ii) for *θ* > 0). The limiting value of 1 for the slope at *θ* = 0 also follows from [24], where it was shown that the pendant edges account for half the expected PD in a pure-birth tree.

Turning to the variance of *R_t_*, the law of total variance gives:

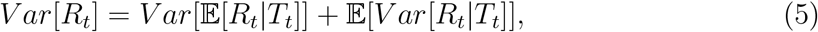

and the first term is zero (since 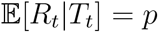, as we have just shown), while the second term is given by:

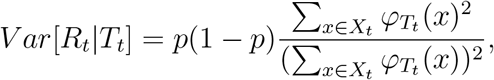

by Eqn (4). Now, for fair proportion and equal splits, we have *φ_T_t__*(*x*) ≤ *t* and therefore:

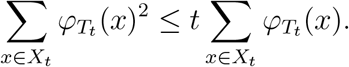

Combining this with Eqn. (5), we obtain

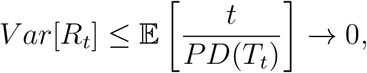

as *t* → ∞ by results from [12], [13] and [22]. Thus *R_t_* converges in probability to 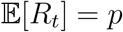. This completes the proof of Lemma 1.

We now state the second technical lemma required (it is a semi-standard result with a straightforward proof that is omitted).

#### Lemma 2.

*Suppose V_t_ and W_t_ are random variables indexed over* 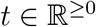 *and suppose that f is a real-valued function that is continuous over the range of W_t_. If W_t_ converges in probability to some constant c, and V_t_ converges in probability to f*(*c*) (*as t* → ∞), *then V_t_* − *f*(*W_t_*) *converges in probability to 0 as t* → ∞.

Returning to the proof of Theorem 1, we now apply Theorem 4.2 (and Corollary 4.3) of [13] to obtain the following result. Let *r* = *b* − *d*, where *b* > *d* > 0. Conditional on the reconstructed tree having at least one leaf, *S_t_*(*p*)/*S_t_*(1) converges in probability to:

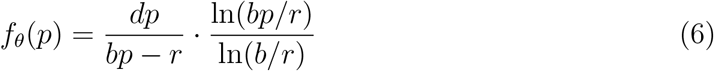

as *t* → ∞ for all values of *r* ≠ *bp* (at this value of *p*, there is no discontinuity in Eqn. (6)). In the case where *d* = 0, (Yule pure-birth), *S_t_*(*p*)/*S_t_*(1) converges in probability to

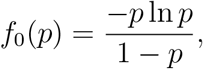

which is also the limiting value of *f_θ_* as *θ* → 0 (i.e. as *d* → 0).

Thus for all *θ* ≥ 0, Δ*PD_T_t__*(*p*)/*PD*(*T_t_*) = 1 − *S_t_*(*p*)/*S_t_*(1) converges in probability to 1 − *f_θ_*(*p*) and, by Lemma 1, Δ*φ_T_t__*(*p*)/*PD*(*T_t_*) converges in probability to 1 − *p* (as *t* → ∞). Thus by Lemma 2:

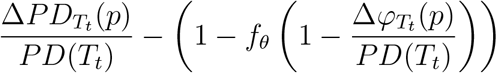

converges in probability to 0 as *t* → ∞. Straightforward (if tedious) algebra now shows that *y_θ_*(*x*) = 1 − *f_θ_*(1 − *x*), and Parts (i) and (ii) now follow. This completes the proof.

## 2. DISCUSSION

The use of evolutionary history as a metric of conservation worth remains a somewhat contentious proposal. This is due, in part, to a bewildering number of possible metrics [28], and confusion regarding what evolutionary history captures: for some researchers, PD captures other values – for example, unmeasured ‘features’ of potential future worth [2, 4] – while for other researchers, PD is valuable *per se* [21, 28]. Here, we deal with the former issue and formalize a connection between two complementary and well-known metrics, Faith’s phylogenetic diversity [2], and a class of species-specific measures of evolutionary isolation [19]. Faith’s PD is a set metric, and was formulated to be used in an explicitly complementary context, for example, to help decide which species or areas should be prioritized based on how much additional PD they would contribute to some growing set of species or places (see [4] for a well worked out example for genera of plants in Southern Africa). Isolation indices such as fair proportion and equal splits, were also created with prioritization of places in mind [18] but were not designed to capture complementarity (D. Redding, pers. comm.). The fair proportion index was, however, subsequently chosen by the Zoological Society of London as a metric to help prioritize at-risk species, with the express intent that the measure captures ‘a species contribution to PD’ [10], such that preserving species that score highly on this measure would conserve more PD. The advantages of species-specific measures were laid out clearly in the papers that first used them: they are attached to species identity, and are simple to use and to communicate.

Both of the isolation measures consider here (fair proportion and equal splits) share certain features: they (i) distribute the entire phylogeny among the leaves such that the sum across the leaves equals the entire PD of the tree in question; (ii) scale with species richness; and (iii) are heavily weighted towards the pendant edge [20]. Therefore, a formal connection between the summed loss of isolation indices and the loss of PD might seem reasonable. Here, we show that when extinction occurs via a ‘field of bullets’ scenario, the relationship is predictable for large phylogenies described by neutral birth–death models (Theorem 1), and the prediction applies well even on medium-sized trees (Figure 2). The key component is the fact that extinction is random with respect to the phylogeny; the examples presented by Faith [3] where isolation does very poorly in capturing PD are examples where extinction was highly clumped on the phylogeny. In this context, it is interesting to note that probabilities of extinction are fully consistent with a field of bullets scenario across birds [11], and, while extinction risk is mildly structured in mammals [6] and amphibians [8], species at higher risk of extinction are not those with higher isolation indices [1, 8], suggesting that summed isolation index loss will be conservative with reference to PD loss. How robust our results are to other patterns of extinction, and whether a field of bullets is the correct model for extinction for other groups, or in other contexts (e.g. when projecting long-term losses of evolutionary history from landscapes due to land conversion) are empirical questions for the future.

## Footnotes

1 Web of Science, accessed May, 2017

